# Dosage-sensitive functions in embryonic development drove the survival of genes on sex-specific chromosomes in snakes, birds, and mammals

**DOI:** 10.1101/2020.07.09.196279

**Authors:** Daniel W. Bellott, David C. Page

**Affiliations:** Whitehead Institute, Cambridge, Massachusetts 02142, USA; Department of Biology, Massachusetts Institute of Technology, Cambridge, Massachusetts 02139, USA; Howard Hughes Medical Institute, Whitehead Institute, Cambridge, Massachusetts 02142, USA

**Author notes:** **Contact** Correspondence should be addressed to David C. Page, Whitehead Institute, 455 Main Street Cambridge, MA 02142 USA.

**Keywords:** comparative genomics, sex chromosomes, haploinsufficiency

## Abstract

Different ancestral autosomes independently evolved into sex chromosomes in snakes, birds, and mammals. In snakes and birds, females are ZW, while males are ZZ; in mammals, females are XX and males are XY. While X and Z chromosomes retain nearly all ancestral genes, sex-specific W and Y chromosomes suffered extensive genetic decay. In both birds and mammals, the genes that survived on sex-specific chromosomes are enriched for broadly expressed, dosage sensitive regulators of gene expression, subject to strong purifying selection. To gain deeper insight into the processes that govern survival on sex-specific chromosomes, we carried out a meta-analysis of survival across 41 species — three snakes, 24 birds and 14 mammals — doubling the number of ancestral genes under investigation and increasing our power to detect enrichments among survivors relative to non-survivors. Out of 2573 ancestral genes, representing an eighth of the ancestral amniote genome, only 322 survive on present-day sex-specific chromosomes. Survivors are enriched for dosage sensitive developmental processes, particularly development of the face. However, there was no enrichment for expression in sex-specific tissues, involvement in sex-determination or gonadogenesis pathways, or conserved sex-biased expression. Broad expression and dosage sensitivity contributed independently to gene survival, suggesting that pleiotropy imposes additional constraints on the evolution of dosage compensation. We propose that maintaining the viability of the heterogametic sex drove gene survival on amniote sex-specific chromosomes, and that subtle modulation of the expression of survivor genes and their autosomal orthologs has disproportionately large effects on development and disease.

## Introduction

Among amniotes, snakes, birds, and mammals are all conspicuous for carrying highly differentiated sex chromosomes. Caenophidian snakes account for roughly 80% of all snake species (Uetz et al. 2019), and unlike boas or pythons, carry cytologically distinguishable sex chromosomes. In caenophidian snakes and birds, females are ZW, while males are ZZ; in therian mammals, females are XX and males are XY (Fig. 1A, Supplemental Fig. S1). None of these three sets of sex chromosomes are orthologous to either of the others (Fridolfsson et al. 1998; Nanda et al. 1999; Ross et al. 2005; Matsubara et al. 2006). Sex-linked genes in one lineage are found on autosomes in the other lineages. For example, genes that are sex-linked in caenophidian snakes are autosomal in chicken (where they reside on chromosomes 2 and 27) and humans (where they are found on chromosomes 3, 7, 10, and 17)(Matsubara et al. 2006). Furthermore, orthologs of sex-linked genes from caenophidian snakes, birds, and mammals are found on separate autosomes in vertebrate outgroup species, like fish (Nakatani et al. 2007; Bellott et al. 2010), indicating that each set of sex chromosomes evolved independently, from what were once ordinary autosomes in the common ancestor of amniotes.

**Figure 1.**
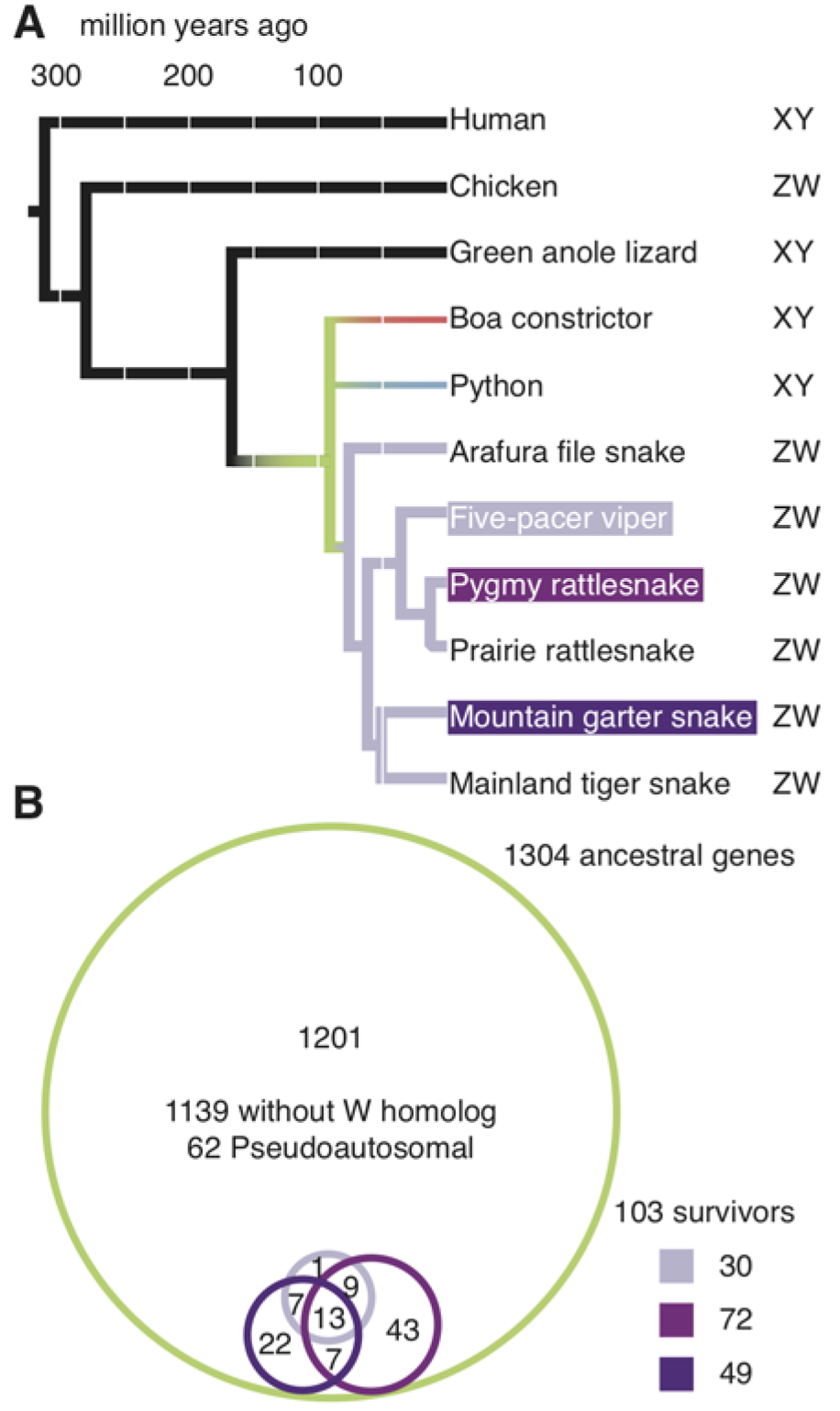
Ancestral Z–W gene pairs from three caenophidian species. (*A*) Phylogenetic tree of selected snake species included in this study, with branches colored to highlight relationships among species. Humans diverged from snakes 312 million years ago. Chicken and Green anole lizard diverged from snakes 280 and 167 million years ago, respectively, and were used to resolve gene gains and losses between snakes and mammals. Snakes diverged from each other starting about 90.8 million years ago (green). Boas (red) and pythons (blue) have independently evolved homomorphic XY sex chromosomes. Caenophidian snakes (purple) share a common ZW sex chromosome system, orthologous to the python XY. (*B*) Euler diagram showing overlapping sets of ancestral Z–W gene pairs identified in five-pacer viper (*light purple*), pygmy rattlesnake (*medium purple*), and mountain garter snake (*dark purple*) as subsets of all 1304 ancestral Z genes (green). See also Supplemental Table S1.

The ancestral autosomes in each of these lineages followed parallel trajectories as they evolved into heteromorphic sex chromosomes. A series of events, most likely inversions on the sex-specific (W or Y) chromosome, gradually suppressed crossing-over between the sex chromosomes, expanding the sex-specific region at the expense of the pseudoautosomal region, and resulting in the formation of evolutionary strata (Lahn and Page 1999; Handley et al. 2004; Schield et al. 2019). In the absence of crossing-over, the sex-specific regions of each new stratum on the W and Y chromosomes began to diverge from their counterparts on the Z and X chromosomes and became subject to genetic decay (Charlesworth and Charlesworth 2000). In birds and mammals, Z and X chromosomes retain greater than 98% of the genes that were present on the ancestral autosomes (Bellott et al. 2010). In contrast, the chicken W chromosome retains only 4% of ancestral genes (Bellott et al. 2017), and the Y chromosomes of therian mammals retain as many as 5% of ancestral genes on the opossum Y chromosome (Bellott et al. 2014), to as few as 1% on the mouse Y chromosome (Bellott et al. 2014).

On the male-specific Y chromosomes of therian mammals, the surviving ancestral genes fall into three functional categories. First, *SRY*, which is responsible for testis determination (Gubbay et al. 1990; Sinclair et al. 1990). Second, testis-expressed genes and gene families retained and amplified to preserve or enhance male fertility (Lahn and Page 1997; Skaletsky et al. 2003; Hughes et al. 2010, 2012; Bellott et al. 2014). Third, broadly-expressed, dosage-sensitive regulators that maintain male viability (Lahn and Page 1997; Bellott et al. 2014).

Analysis of W-linked genes across chickens and 13 additional avian species showed that survival of ancestral genes was biased towards preserving dosage-sensitive regulators expressed across many tissues. However, in contrast to mammals, all ancestral genes in the chicken are broadly expressed, suggesting a focus on preserving female viability (Bellott et al. 2017).

Based on cytological observations of caenophidian snakes, all snakes had been assumed to share a common ZW sex determination system (Ohno 1967; Vicoso et al. 2013a), but the recent discovery of male-specific sequences in *Boa constrictor, Boa imperator*, and Burmese python (*Python bivittatus*) indicated that boas and pythons independently evolved X and Y sex chromosomes (Gamble et al. 2017) (Fig. 1A). Caenophidian snakes share a common ZW system that evolved after their divergence from boas and pythons, but before the divergence of wart and file snakes from other caenophidians (Rovatsos et al. 2015; Matsubara et al. 2016), between 91 and 76 million years ago (Kumar et al. 2017) (Fig. 1A). Comparisons of male and female whole-genome shotgun sequence data from five-pacer viper (*Deinagkistrodon acutus*)(Yin et al. 2016), pygmy rattlesnake (*Sistrurus miliarius*) and mountain garter snake (*Thamnophis elegans*)(Vicoso et al. 2013a) yielded dozens of W-linked gene predictions from each species. Despite being evolutionarily younger than the W chromosomes of birds (greater than 111 million years) or the Y chromosomes of therian mammals (greater than 159 million years)(Kumar et al. 2017), genetic decay is estimated to have been similarly extensive; only 3% of ancestral genes were retained on the W chromosome of the five-pacer viper (Yin et al. 2016). The recently published chromosome-level genome assembly of the prairie rattlesnake (*Crotalus viridis*) includes a Z chromosome that contains around 1500 genes, 30% more than were included in reconstructions based on whole-genome shotgun data (Schield et al. 2019), and more genes than were present on the ancestral autosomes that became the sex chromosomes of birds and therian mammals, combined.

Thus, the sex chromosomes of caenophidian snakes provide a third and much more extensive replicate in a natural experiment where otherwise ordinary autosomes are exposed to genetic decay and the evolutionary pressures that accompany sex chromosome evolution.

We sought to examine whether evolutionary pressures analogous to those we characterized on the male-specific Y chromosomes of therian mammals and the female-specific W chromosomes of birds also shaped the gene content of the caenophidian W chromosome. Further, we wondered whether we might detect more subtle influences on the survival of ancestral genes by combining data from these three lineages. Additionally, we looked for evidence that genes on the ancestral autosomes with functions related to sex and reproduction were more likely to survive on sex-specific chromosomes.

Here we reconstruct the ancestral set of genes on the caenophidian sex chromosomes and trace their evolutionary trajectory. We show that the survivors of genetic decay in caenophidian snakes, like the survivors in birds and mammals, are enriched for broadly-expressed, dosage sensitive regulators evolving under strong purifying selection. We combined the ancestral genes from all three lineages to analyze the evolutionary pressures influencing survival as genetic decay winnowed 2573 ancestral genes (representing an eighth of the ancestral amniote genome) down to 322 survivors through 2.5 billion years of sex chromosome evolution. We find that factors related to sex and reproduction had little influence on the survival of ancestral genes on sex-specific chromosomes; survivors are not enriched for involvement in sex-determination or gonadogenesis pathways, for conserved expression in sex-specific tissues, or for conserved sex-biased expression. However, dosage sensitivity and breadth of expression made strong and independent contributions to the survival of genes on sex-specific chromosomes. We conclude that, in amniotes, selection to maintain the viability of the heterogametic sex and preserve the ancestral dosage of broadly-expressed homologous Z–W and X–Y gene pairs was the driving force behind survival in the face of widespread genetic decay on sex-specific W and Y chromosomes.

## Results

### Reconstructing the ancestral autosomes

To analyze the caenophidian sex chromosomes for biases in gene content that arose during sex chromosome evolution, we reconstructed the gene content of the ancestral autosomes from which they evolved. Previous reconstructions of caenophidian sex chromosome evolution used lizard chromosome 6 as a proxy for the ancestral autosomes, and identified 1135 ancestral Z genes (Vicoso et al 2013a, Yin et al 2016). Beginning with genes previously identified as sex-linked in snakes (Supplemental Table S1), we searched for syntenic orthologs in the assemblies of three caenophidian snakes — prairie rattlesnake (Schield et al. 2019), mainland tiger snake (*Notechis scutatus*)(Edwards et al. 2020), and five-pacer viper (Yin et al. 2016) — using orthologous autosomal scaffolds from the assembly of the *Boa constrictor* (Bradnam et al. 2013) as an out-group to resolve lineage-specific gains and losses within snakes (Fig. 1A). We identified 45% more genes than previous reconstructions: a total of 1652 genes in 1304 protein-coding gene families on the ancestral autosomes that became the caenophidian Z and W sex chromosomes (Fig. 1B, Supplemental Table S2). This set of ancestral genes is comprehensive, extending across the Z chromosome to include 62 single-copy genes in the pseudoautosomal region (Fig. 1B, Supplemental Table S2, Supplemental Figs. S2 and S3). We restricted our subsequent analyses of the decay and survival of ancestral genes to the 1242 protein-coding gene families outside the pseudoautosomal region.

We identified survivors among ancestral genes by comparing our list of ancestral genes to published W-linked genes from five-pacer viper (Yin et al. 2016), pygmy rattlesnake, and mountain garter snake (Vicoso et al. 2013a) (Supplemental Table S3), including candidate W-linked genes previously excluded on the basis of their chromosomal assignment in green anole lizard (Supplemental Table S4). We identified 103 ancestral genes with surviving W homologs in one of these three species (Fig. 1B, Supplemental Table S2). We compared these 103 surviving ancestral Z–W gene pairs to the other 1139 ancestral genes without surviving W homologs (Fig. 1B). Using published data sets, we searched for evidence that Z–W pair genes systematically differ from ancestral genes on the Z chromosome that lack W homologs with regard to dosage sensitivity (Huang et al. 2010; Ruderfer et al. 2016; Agarwal et al. 2015), breadth of expression (Perry et al. 2018; Merkin et al. 2012; Ayers et al. 2013; Petropoulos et al. 2016), and intensity of purifying selection (Zerbino et al. 2018) (Fig. 2, Supplemental Table S5).

**Figure 2.**
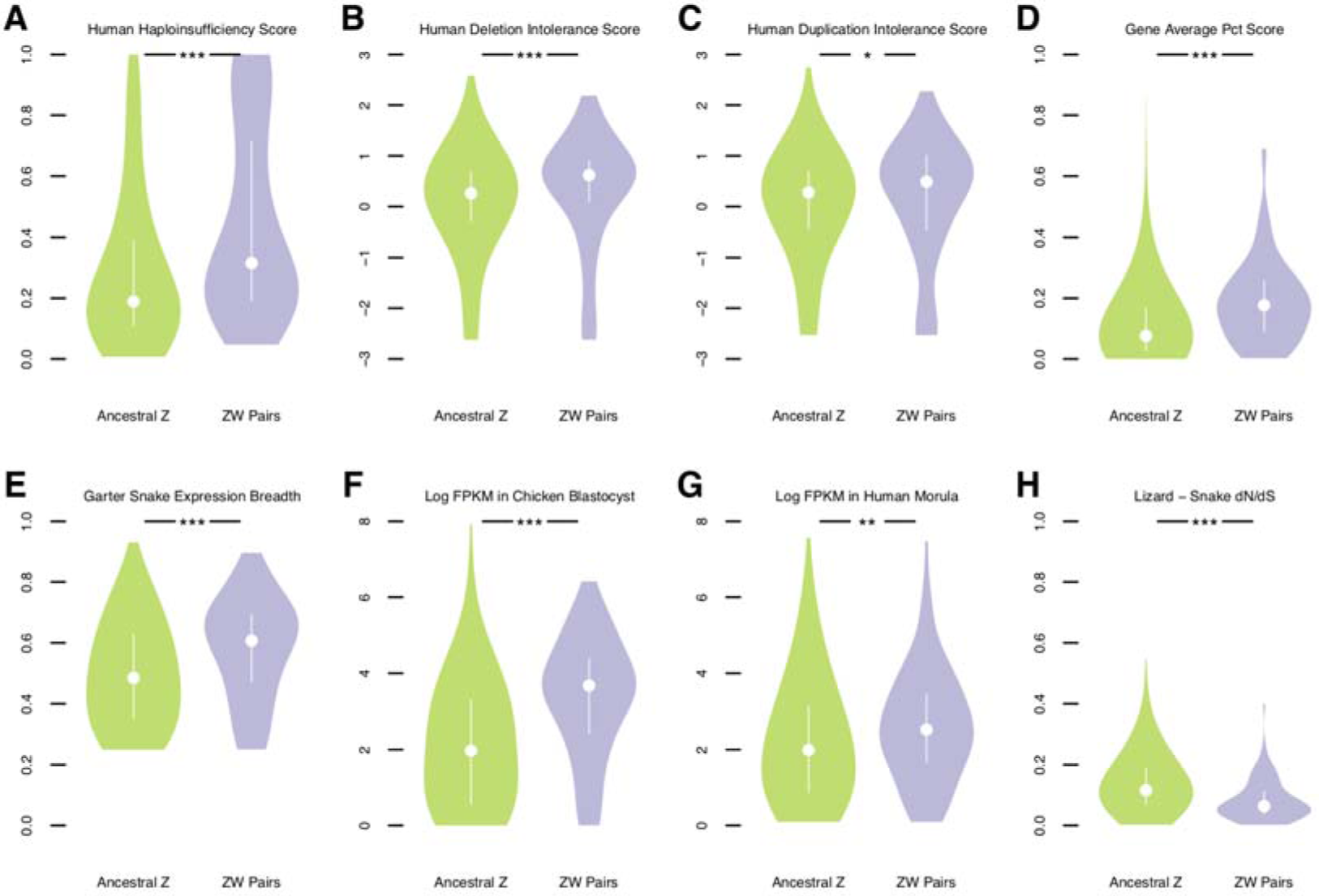
Factors in the survival of caenophidian Z–W gene pairs. Violin plots, with the median (white circle) and interquartile range (white bar) indicated, compare annotations of ancestral Z–W gene pairs identified in 3 species (purple) to annotations for the remainder of ancestral Z genes (green). (*) P < 0.05, (**) P < 0.01, (***) P < 0.001. P values obtained using one-tailed Mann–Whitney U tests. See Methods and Supplemental Table S15. Human orthologs of ancestral caenophidian Z–W gene pairs have greater (*A*) probability of haploinsufficiency, (*B*) deletion intolerance scores, (*C*) duplication intolerance scores, and (*D*) mean probabilities of conserved targeting (PCT) than other ancestral Z genes. Orthologs of ancestral Z–W gene pairs are more broadly expressed than orthologs of other ancestral Z genes (*E*) in a panel of seven adult eastern garter snake tissues. Orthologs of ancestral Z–W gene pairs are more highly expressed than orthologs of other ancestral Z genes (*F*) in chicken blastocysts and (*G*) in human preimplantaion embryos. Orthologs of ancestral Z–W gene pairs have reduced dN/dS ratios compared to orthologs of other ancestral Z genes in alignments between (*H*) tiger snake and green anole lizard orthologs.

### Caenophidian Z–W gene pairs are dosage sensitive

We looked for signs of dosage sensitivity among surviving caenophidian Z–W gene pairs. We mapped various published gene-by-gene measures of dosage sensitivity from the human genome onto their snake orthologs (Supplemental Table 5). Using gene-by-gene estimates of the probability of haploinsufficiency in humans (Huang et al. 2010), we found that human orthologs of surviving Z–W gene pairs were more likely to be haploinsufficient than the human orthologs of ancestral Z-chromosome genes that lack W-chromosome homologs (Fig. 2A). Likewise, using deletion intolerance scores calculated from a catalog of genic copy number variation among 59,898 control human exomes (Exome Aggregation Consortium [ExAC])(Ruderfer et al. 2016), we found that human orthologs of surviving Z–W gene pairs were less likely to tolerate deletions than the orthologs of ancestral Z-chromosome genes that lack W-chromosome homologs (Fig. 2B).

We previously observed an enrichment for human disorders caused by haploinsufficiency among the orthologs of chicken Z–W gene pairs, relative to other ancestral genes. Of the 103 caenophidian Z–W gene pairs, at least 11 have human orthologs implicated in congenital disorders caused by heterozygous loss-of-function mutations (Table 1). We assessed the likelihood of finding 11 haploinsufficient human orthologs among these 103 Z–W pairs by examining all 299 human orthologs of ancestral caenophidian Z-linked genes linked to a phenotype with a known molecular basis in Online Mendelian Inheritance in Man (OMIM)(McKusick-Nathans Institute of Genetic Medicine 2019)(Table 2). We found that Z–W pair genes were enriched three-fold for phenotypes due to haploinsufficiency relative to Z-linked genes without surviving W-homologs. Taken together, the elevated haploinsufficiency probabilities, deletion intolerance scores, and significant enrichment for human disorders caused by haploinsufficiency indicate that caenophidian Z–W pairs are enriched for haploinsufficient genes.

**Table 1.**
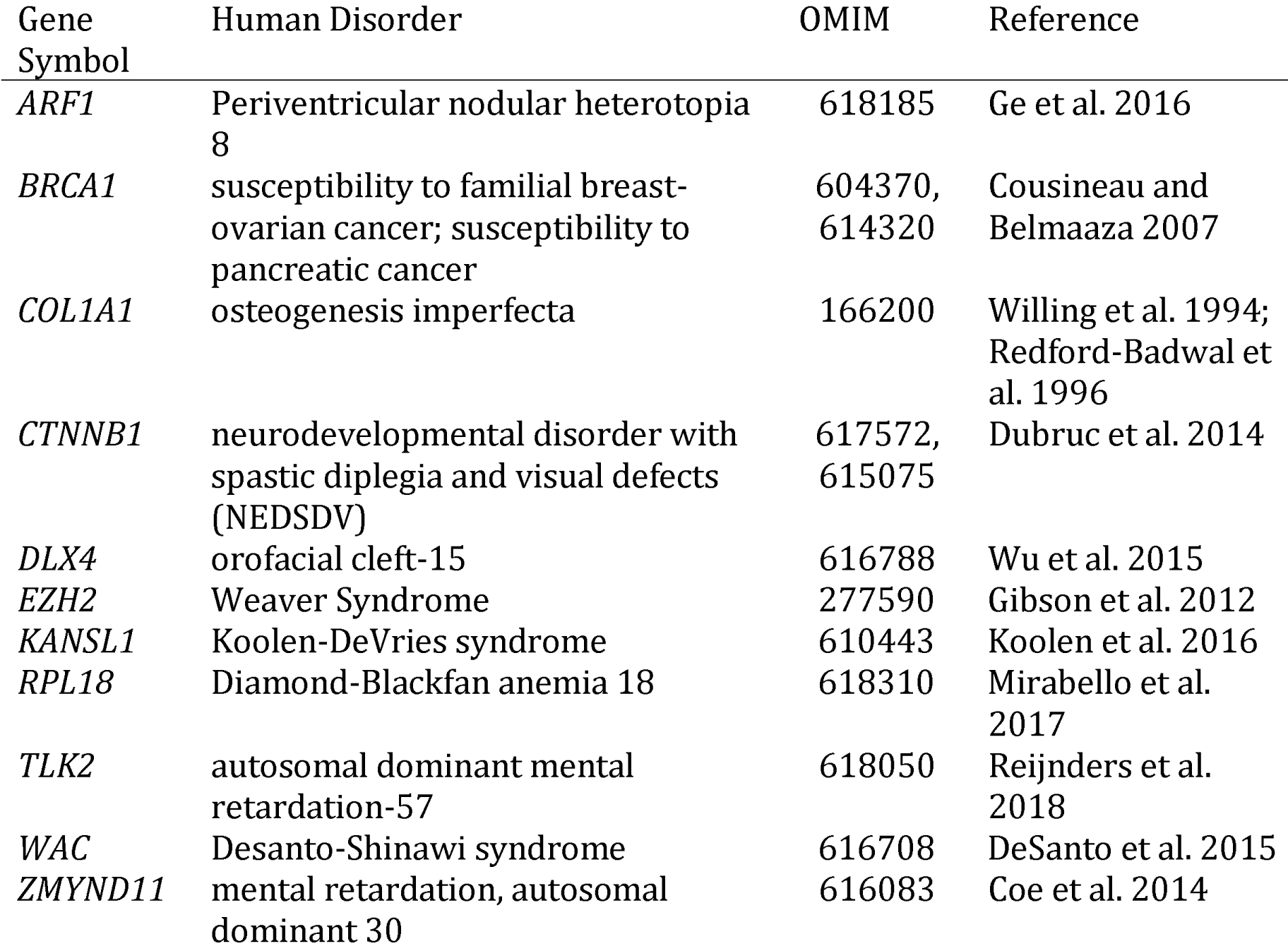
Human orthologs of caenophidian Z–W pairs linked to human disorders attributed to haploinsufficiency.

| Gene Symbol | Human Disorder | OMIM | Reference |
| --- | --- | --- | --- |
| *ARF1* | Periventricular nodular heterotopia 8 | 618185 | Ge et al. 2016 |
| *BRCA1* | susceptibility to familial breast-ovarian cancer; susceptibility to pancreatic cancer | 604370, 614320 | Cousineau and Belmaaza 2007 |
| *COL1A1* | osteogenesis imperfecta | 166200 | Willing et al. 1994; Redford-Badwal et al. 1996 |
| *CTNNB1* | neurodevelopmental disorder with spastic diplegia and visual defects (NEDSDV) | 617572, 615075 | Dubruc et al. 2014 |
| *DLX4* | orofacial cleft-15 | 616788 | Wu et al. 2015 |
| *EZH2* | Weaver Syndrome | 277590 | Gibson et al. 2012 |
| *KANSL1* | Koolen-DeVries syndrome | 610443 | Koolen et al. 2016 |
| *RPL18* | Diamond-Blackfan anemia 18 | 618310 | Mirabello et al. 2017 |
| *TLK2* | autosomal dominant mental retardation-57 | 618050 | Reijnders et al. 2018 |
| *WAC* | Desanto-Shinawi syndrome | 616708 | DeSanto et al. 2015 |
| *ZMYND11* | mental retardation, autosomal dominant 30 | 616083 | Coe et al. 2014 |

**Table 2.** Human Disorders in OMIM attributed to haploinsufficiency.

|  | Haploinsufficient | Not Haploinsufficient |
| --- | --- | --- |
| Z–W Pairs | 11 | 18 |
| Other Z | 34 | 236 |

We previously reported that, in addition to being sensitive to loss of gene dosage, therian X–Y and avian Z–W gene pairs are more sensitive to over-expression as well (Naqvi et al. 2018). Using the EXAC duplication intolerance scores (Ruderfer et al. 2016), we find that, relative to other ancestral Z genes, the human orthologs of caenophidian Z–W gene pairs are more intolerant of duplication (Fig. 2C). MicroRNAs (miRNAs) are short, noncoding RNAs that fine-tune gene dosage by pairing to the 3’ untranslated region and lowering target mRNA levels (Bartel 2009). Conserved targeting by miRNAs is a feature of genes sensitive to changes in gene dosage, particularly to over-expression (Naqvi et al. 2018). We used published estimates of the probabilities of conserved targeting (PCT scores)(Friedman et al. 2009) for each gene-miRNA interaction in the human genome. We found that the human orthologs of surviving Z–W gene pairs had higher gene-average PCT scores than other ancestral Z genes (Fig. 2D). Also, across all gene-miRNA pairs, PCT scores are higher for Z–W gene pairs than for other ancestral Z genes (Supplemental Fig. S4A). Moreover, relative to other ancestral genes, the orthologs of Z–W gene pairs are targeted by more miRNA sites conserved between human and chicken (Supplemental Fig. S4B), and between human and green anole lizard (Supplemental Fig. S4C). We conclude that caenophidian Z–W pairs are similar to both therian X–Y and avian Z–W gene pairs in being more sensitive to increases and decreases in gene expression than other ancestral genes.

### Caenophidian Z–W gene pairs are broadly expressed

We previously observed that, relative to other genes that were present on the ancestral autosomes, therian X–Y and avian Z–W gene pairs are enriched for genes broadly expressed across adult tissues and through developmental time (Bellott et al. 2014, 2017). We assessed whether surviving caenophidian Z–W pairs are more broadly expressed than other ancestral genes in snakes, and whether this broad expression was ancestral. We observed that the Z homologs of Z–W gene pairs are more broadly expressed than ancestral genes that lack W homologs across a panel of adult eastern garter snake (*Thamnophis sirtalis*) female tissues (Perry et al. 2018) (Fig. 2E). This increased breadth of expression extends to chicken (Supplemental Fig. S4D) and human (Supplemental Fig. S4E) orthologs of Z–W gene pairs. We conclude that caenophidian Z–W gene pairs maintain broad expression across adult tissues from their autosomal precursors in the amniote ancestor.

We further suggest that this breadth of expression extended to the earliest stages of embryonic development. While gene expression data from snake embryos is not available, we found that the orthologs of ancestral caenophidian Z–W gene pairs are more highly expressed than those of ancestral Z genes that lack W homologs in both chicken blastocysts (Ayers et al. 2013) (Fig. 2F) and human morulae (Petropoulos et al. 2016) (Fig. 2G). We conclude that the autosomal precursors of caenophidian Z–W pairs were more broadly expressed across development as well as across adult tissues.

### Caenophidian Z–W gene pairs are subject to strong purifying selection

Previously we observed that the X-homologs of therian X–Y pairs (Bellott et al. 2014), and the Z-homologs of avian Z–W pairs (Bellott et al. 2017), are under stronger purifying selection than ancestral genes without Y or W homologs. If surviving X–Y and Z–W gene pairs are extremely dosage sensitive, alleles that alter the ancestral function of the homologs on the X or the Z chromosomes should be detrimental to both males and females.

We examined Ensembl mainland tiger snake alignments (Zerbino et al. 2018) for evidence that the Z-linked homologs of caenophidian Z–W gene pairs are subject to stronger purifying selection than other ancestral Z-linked genes. Compared to ancestral genes on the caenophidian Z chromosome that lack W homologs, the Z homologs of Z–W gene pairs have a reduced ratio of non-synonymous to synonymous substitution (dN/dS) rates when mainland tiger snake genes are compared to their orthologs in green anole lizard (Fig. 2H), chicken (Supplemental Fig. S4F), and human (Supplemental Fig. S4G). We conclude that, on caenophidian sex chromosomes, strong purifying selection preserved a subset of ancestral genes with pre-existing broad expression and dosage sensitivity, dating back to the common ancestor of amniotes.

### Functional coherence of caenophidian Z–W gene pairs

Human X–Y gene pairs and chicken Z–W gene pairs perform regulatory functions in key cellular processes, positioning them to govern the expression of targets throughout the genome (Bellott et al. 2014, 2017). In both lineages, survivors play roles in transcription, translation, and protein stability, and while human X–Y gene pairs have a greater emphasis on mRNA splicing, many chicken Z–W pairs act in protein secretion and signal transduction pathways (Bellott et al. 2017).

We asked whether the set of ancestral genes that survived on caenophidian W chromosomes carried out regulatory functions that parallel those of mammals and birds. In comparison to other ancestral genes on the caenophidian Z chromosome, Z–W pair genes are enriched for Gene Ontology (GO) annotations such as nucleic acid binding, nucleus, and transcription (Supplemental Table S6) that suggest regulatory functions. We looked in more detail at the molecular functions of the 103 caenophidian Z–W pairs (Fig. 3)(McKusick-Nathans Institute of Genetic Medicine 2019; The UniProt Consortium 2019). We observe that caenophidian Z–W pairs share functions in regulating transcription and protein stability with surviving gene pairs in human and chicken (Fig. 3). Four survivors are components of the proteasome (*PSMA2, PSME3, PSMD3*, and *PSMD11*) (Fig. 3), consistent with the idea that members of large protein complexes tend to be dosage-sensitive (Papp et al. 2003; Pessia et al. 2012).

**Figure 3.**
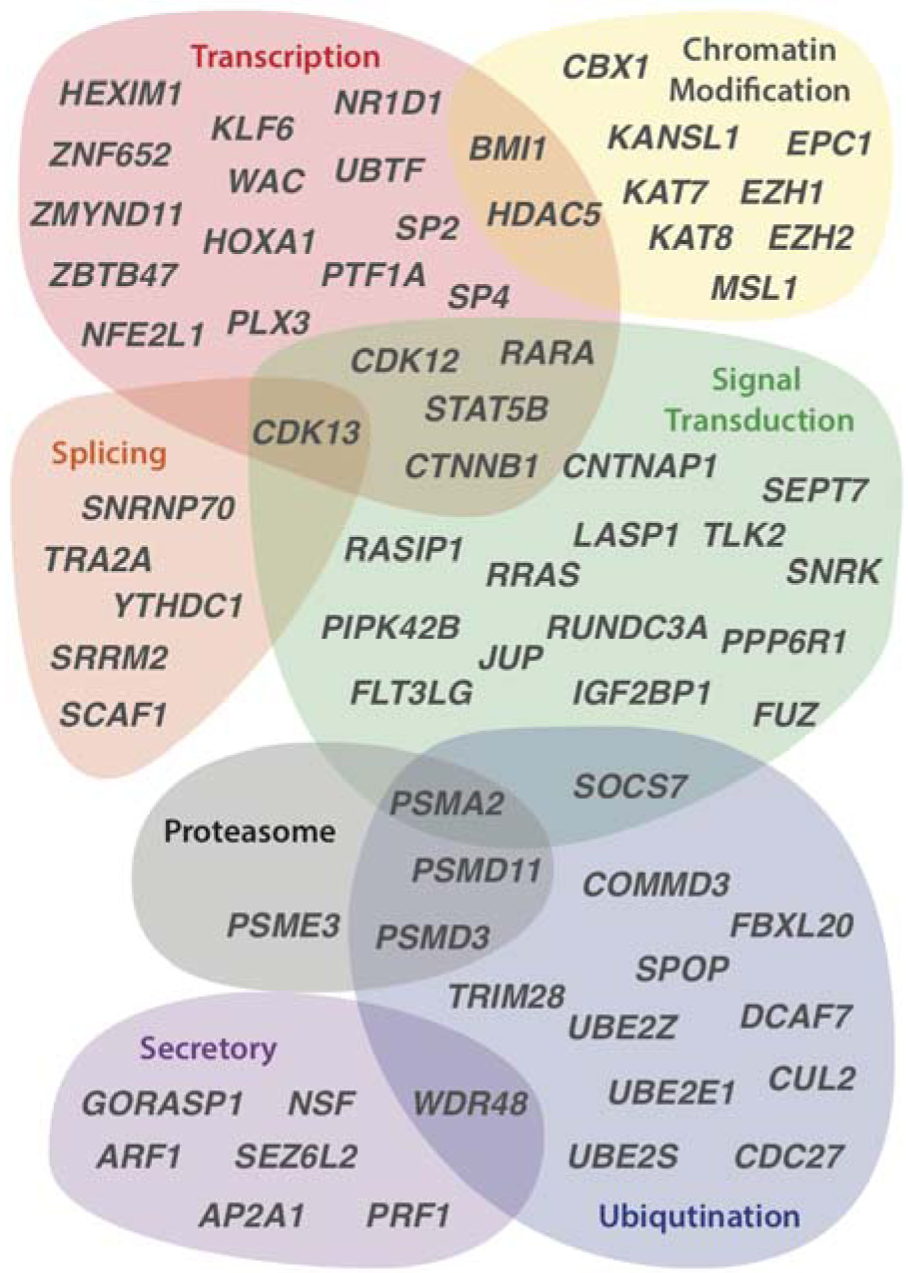
Regulatory annotations of ancestral caenophidian Z–W gene pairs. The Euler diagram depicts regulatory functions predicted for genes from selected Z– W gene pairs on the basis of UniProt annotations of human orthologs. See also Supplemental Table S6.

The annotations of caenophidian Z–W pairs also indicate overlap with functions we previously observed only in mammals or chicken; as in human, they play roles in regulating mRNA splicing, and, similar to chicken, they participate in secretory and signal transduction pathways. We previously observed that chicken Z–W pairs included several components of the Wnt signaling pathway; we note that both junction plackoglobin (*JUP*) and β-catenin (*CTNNB1*) are Z–W pairs in caenophidian snakes. This underscores the importance of the correct gene dose for proper signaling in pathways regulating growth and development.

### Pressure to preserve dosage-sensitive functions in early embryonic development drove gene survival

Since our analyses of snakes revealed broad functional categories similar to those we previously observed in birds and mammals, we reasoned that incorporating all three lineages into a combined analysis might give us the power to detect more specific functional enrichments than we recovered from individual lineages. We therefore combined our data on the survival of ancestral genes across all three lineages, and incorporated published survivors from additional birds (Xu et al. 2019a) and mammals (Li et al. 2013; Skinner et al. 2016; Janečka et al. 2018). From 41 species — three snakes, 24 birds, and 14 mammals — we compiled a list of 322 survivors of genetic decay from a total of 2573 ancestral genes (Supplemental Fig. S1, Supplemental Table S5), representing nearly an eighth of the ancestral amniote genome. We first verified that combining our data did not eliminate or reverse the enrichments we previously observed; the combined set of surviving ancestral genes is enriched for the same qualities we observed separately in each lineage (Supplemental Fig. S5). Therefore, survivors are enriched for the same broad categories of dosage-sensitive, broadly-expressed genes evolving under strong purifying selection across all three amniote lineages.

This broad sampling of survivors allowed us to detect more specific enrichments, providing insight into the developmental processes evolving under strong purifying selection across amniotes. Relative to non-survivors, survivors were enriched for GO terms that emphasize their role in regulating development: “developmental growth”, “chordate embryonic development”, “regulation of anatomical structure morphogenesis”, “regulation of epithelial to mesenchymal transition”, “renal system development”, and “face development” (Supplemental Table S7).

These annotations suggest that pressure to preserve dosage-sensitive functions in early embryonic development drives the survival of ancestral genes on sex-specific chromosomes. The epithelial to mesenchymal transition is critical to gastrulation; haploinsufficiency for genes governing this process could explain why most 45,X human conceptuses spontaneously abort in the first trimester with an absent or disorganized embryo (Cockwell et al. 1991; Hook and Warburton 1983), and why chicken embryos carrying a single Z chromosome are observed at the blastocyst stage (Fechheimer 1981), but not at 4-5 days of development (Bloom 1972).

Shortly after gastrulation, neural crest cells undertake an epithelial to mesenchymal transition, migrate through the embryo, and differentiate into a diverse array of tissues, including the bones and cartilage of the face. Neural crest cells respond to a multitude of signal-transduction pathways, and they are sensitized to p53-mediated apoptosis (Calo, et al. 2018). As a result, mutations in hundreds of genes can cause defects in the specification, migration, differentiation, and death of neural crest cells, resulting in a wide range of pathologies, including craniofacial anomalies (Trainor, 2010; Vega-Lopez, et al. 2018). We conclude that the enrichment for “face development” reflects the extreme sensitivity of the neural crest to changes in the dose of broadly expressed survivor genes, rather than an enrichment for face-specific expression.

The face represents a highly sensitive assay for disruptions to early human development throughout the body. The enrichment for minor physical anomalies, particularly of the head and face, in disorders of the central nervous system, like autism, intellectual disability, schizophrenia, and bipolar disorder, has been taken as evidence that these disorders are developmental in origin (Smith and Bostian 1964, Weinberg et al. 2007, Aldridge et al. 2011, Berecz et al. 2017). The sets of genes implicated in neurodevelopmental disorders overlap with each other and share characteristics with genes that survive on sex-specific chromosomes: an enrichment for chromatin modifiers and embryonic expression, as well as signs of haploinsufficiency (Iossifov et al. 2014). We therefore looked for evidence that the human orthologs of survivor genes were enriched among a set of 1919 genes identified as carrying *de novo* mutations in whole exome sequencing studies across two or more neurodevelopmental disorders, including autism spectrum disorders, bipolar disorder, intellectual disability, or schizophrenia (Wang, Corominas, and Lin 2019)(Table 3, Supplemental Table S5). We found that orthologs of survivors were enriched 1.6-fold for genes mutated in neurodevelopmental disorders compared with non-survivors. The genes implicated in neurodevelopmental disorders overlap with those implicated in congenital anomalies of other organ systems, for example, those of the heart or the kidney and urinary tract (Homsy et al. 2015, Sanna-Cherchi et al. 2018). Because survivor genes are broadly expressed, we asked whether they were also enriched among genes implicated in these disorders, using a set of 1169 genes associated with congenital heart defects (Homsy et al. 2015)(Table 4, Supplemental Table S5) and 50 genes associated with congenital anomalies of the kidney and urinary tract (Sanna-Cherchi et al. 2018)(Table 5, Supplemental Table S5). We found that human orthologs of survivors were enriched 1.6-fold for genes mutated in congenital heart defects compared to non-survivors (Table 4). Likewise, human orthologs of survivors were enriched four-fold for genes associated with congenital anomalies of the kidney and urinary tract compared to non-survivors (Table 5). We conclude that the early developmental role of survivors may have far-reaching consequences for fitness through their effects not only on the face, but also the brain, heart, and kidney. Close scrutiny of the human orthologs of survivors in snakes and birds will likely reveal new candidates for human disorders and insights into the developmental pathways most sensitive to changes in gene dose.

**Table 3.** Survivors enriched in neurodevelopmental disorders.

|  | Mutated in Neurodevelopmental Disorders | Not Mutated in Neurodevelopmental Disorders |
| --- | --- | --- |
| Survivors | 48 | 274 |
| Non-Survivors | 205 | 2046 |

**Table 4.** Survivors enriched in congenital heart disease.

|  | Mutated in Neurodevelopmental Disorders | Not Mutated in Neurodevelopmental Disorders |
| --- | --- | --- |
| Survivors | 28 | 294 |
| Non-Survivors | 121 | 2130 |

**Table 5.** Survivors enriched in congenital anomalies of the kidney and urinary tract.

|  | Mutated in Neurodevelopmental Disorders | Not Mutated in Neurodevelopmental Disorders |
| --- | --- | --- |
| Survivors | 4 | 318 |
| Non-Survivors | 6 | 2245 |

Combining the data from all three lineages also revealed a nine-fold enrichment for components of the ficolin-1-rich granule lumen (Supplemental Table S7), a cellular compartment that may play an under-appreciated role in regulating neural crest development. There were 12 survivors out of 22 ancestral genes in this GO category, including survivors from all three lineages (Table 6). Relative to the rest of the genome, genes annotated with this GO category are more likely to be haploinsufficient (Supplemental Fig. S6). Ficolin-1 is a recognition molecule in the lectin complement pathway of the innate immune system, localized in secretory granules in the cytoplasm of neutrophils and monocytes (Liu et al. 2005). Apart from their role in innate immunity, components of the complement cascade are involved in pruning inappropriate synapses during development of the central nervous system (Stevens et al. 2007), and mutations in components upstream of the complement cascade are responsible for 3MC syndrome (also known as Craniofacial-Ulnar-Renal syndrome or Carnevale, Mingarelli, Malpuech and Michels syndromes) in humans, as a result of their role in guiding neural crest cell migration (Rooryck et al. 2011; Munye et al. 2017). Several survivors annotated ficolin-1-rich granule components — *CDK13, DDX3X, HUWE1*, and *JUP* — are implicated in human disorders with dysmorphic facial features, cleft lip and palate, poor growth, intellectual disability, hearing loss, and cardiac anomalies that suggest a role in regulating neural crest development. We speculate that regulation of face and renal development by the complement pathway through ficolin-1-rich granules is a dosage-sensitive process, like the TGF-β and Wnt pathways that are also enriched among surviving Z–W pairs in birds and snakes.

**Table 6.**
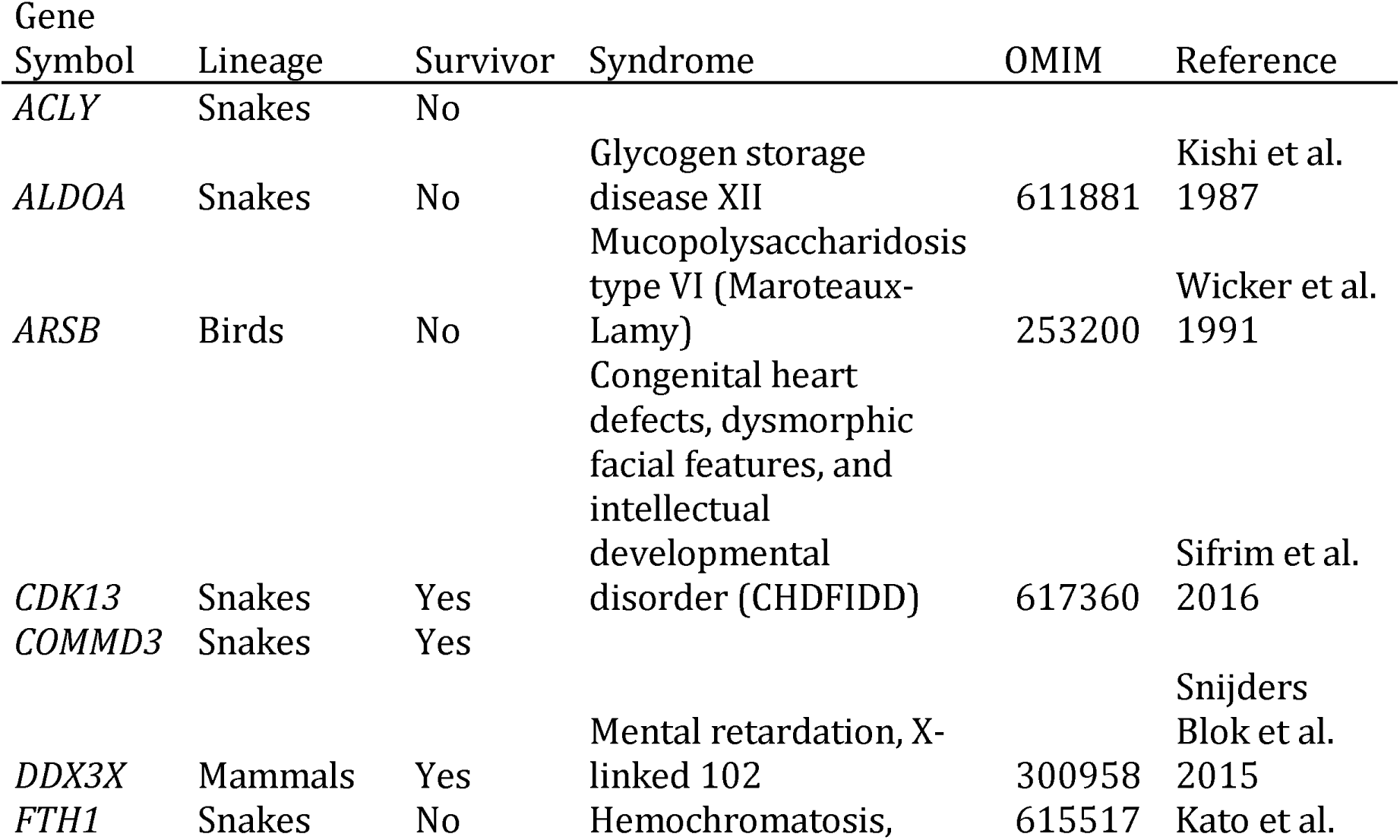
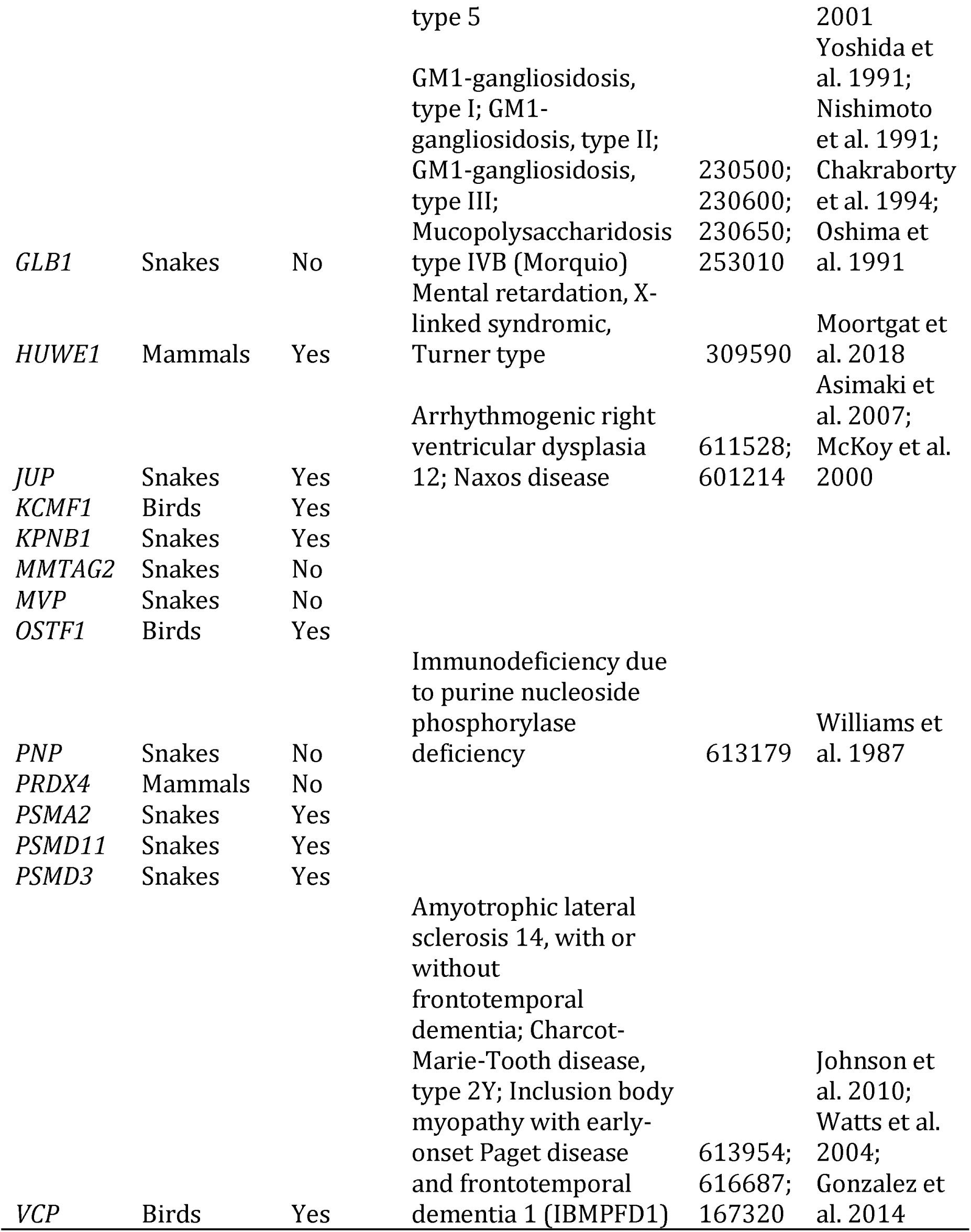
Ficolin-1-rich granule lumen components among ancestral genes.

### Sex determination, sex specific expression, and gene survival on sex chromosomes

Across all three lineages, ancestral X–Y and Z–W gene pairs were preserved to maintain comparable expression between males and females, but other evolutionary strategies also contribute to gene survival on the male-specific Y chromosome of mammals. The survival of *SRY* is founded on its unique status as the testis-determining gene (Gubbay et al. 1990; Sinclair et al. 1990), while selective pressures to preserve or enhance male fertility drove the retention and amplification of testis-expressed gene families (Lahn and Page 1997, Bellott et al. 2014). Analogous strategies are predicted to act on W chromosomes, causing the accumulation of genes expressed predominantly in female-specific tissues (Moghadam et al. 2012; Mank et al. 2014). There is little evidence that these strategies operate in chicken, where the identity of the master sex determining gene is unknown (Zhao et al. 2010), and all surviving genes on the W chromosome, including the highly amplified *HINTW* gene family, are broadly expressed in females (Ayers et al. 2013; Bellott et al. 2017).

Nevertheless, sex-determining genes (those involved in the sex determination cascade and gonadogenesis) and genes with sex-biased functions (those with expression limited to sex-specific tissues, or with sex-biased gene expression in somatic tissues) play a large role in theoretical models of sex chromosome evolution. Linkage between sex-determining genes and genes with sex-biased functions is predicted to drive the suppression of recombination on sex chromosomes (Nei 1969; Bull 1983; Rice 1987), resulting in the formation of evolutionary strata and heteromorphic sex chromosomes. This has led to speculation that certain ancestral amniote chromosomes may have been predisposed to take on roles as sex chromosomes, due to an enrichment for genes in the sex-determination pathway linked to other genes with sex-biased functions (O’Meally et al. 2012).

We looked for evidence that ancestral sex and reproduction related functions promoted the survival of Y and W-linked genes across amniotes. We reanalyzed published data sets for evidence that survivors systematically differ from other ancestral genes with regard to the prevalence of genes involved in sex determination and gonadogenesis (Ayers et al. 2013), gene expression limited to sex-specific tissues (GTEx Consortium 2017; McCarthy et al. 2019; Perry et al. 2018; Yin et al. 2016), and conserved sex-biased expression (Naqvi et al. 2019) (Fig. 4, Tables 2 and 3, Supplemental Table S5). We were unable to detect strong associations between any of these categories and gene survival on sex-specific chromosomes.

**Figure 4.**
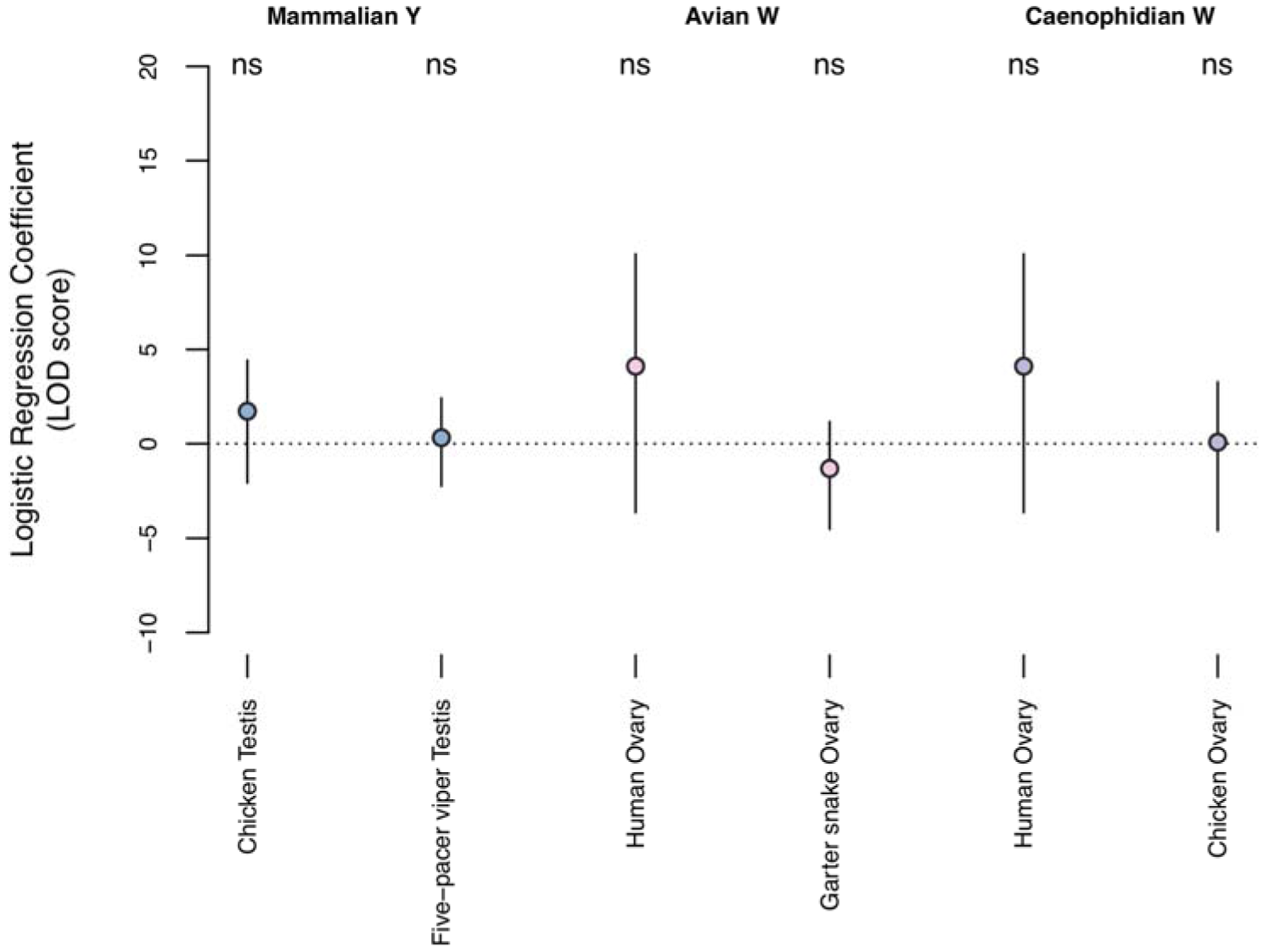
Survivors are not ancestrally specialized for expression in reproductive tract tissues. Coefficients for testis- and ovary-specific expression (circles) and 95% confidence intervals (bars), from logistic regression models of gene survival on sex-specific chromosomes in mammals (blue), birds (pink) and snakes (purple), conditional on breadth of expression. (ns) P > 0.05. After controlling for expression breadth, the regression coefficient is not significantly different from 0 for any tissue from any species.

### Survivors are not significantly enriched for sexual reproduction and gonadogenesis genes

We asked whether ancestral genes with a surviving Y or W-linked homolog were more likely than other ancestral genes to be present in a list of 117 genes implicated in sexual reproduction or gonadogenesis in vertebrate model organisms (Ayers et al. 2013). We did not detect a significant enrichment for genes involved in sexual reproduction or gonadogenesis among survivors; while 7 of 324 survivors were on the list, so were 28 of 2249 non-survivors (Table 7).

**Table 7.**
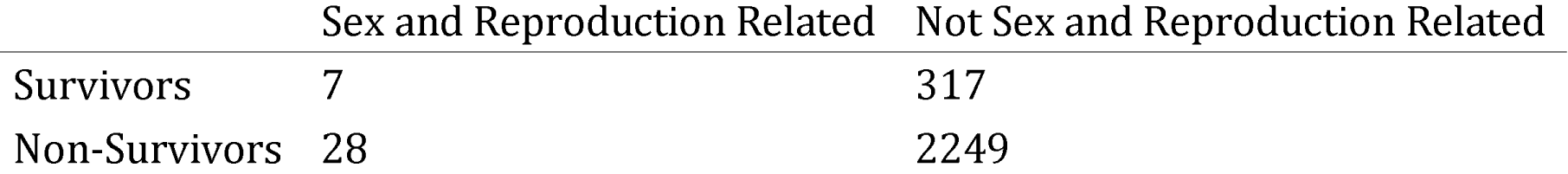
Survival of sex and reproduction related genes.

|  | Sex and Reproduction Related | Not Sex and Reproduction Related |
| --- | --- | --- |
| Survivors | 7 | 317 |
| Non-Survivors | 28 | 2249 |

### Survivors are not significantly enriched for genes highly expressed in sex-specific tissues

We reasoned that genes with ancestrally sex-biased functions might be expressed more highly in sex-specific tissues, including those of the reproductive tract, and therefore we looked for evidence that the survivors of genetic decay are enriched for tissue specific expression in adult gonads in human, chicken, and snakes (GTEx Consortium 2017; McCarthy et al. 2019; Perry et al. 2018; Yin et al. 2016). For each gene-tissue pair, we calculated a tissue specificity index, dividing the expression in a tissue by the summed gene expression across all tissues (Supplemental Table S5). Most survivors showed only weak specificity in sex-specific tissues, consistent with their being expressed at nearly equal levels across adult tissues (Supplemental Table S5). Therefore, we used logistic regression to control for the effects of expression breadth while measuring the effect of tissue specific expression on survival. We analyzed each lineage separately, using the expression patterns of the autosomal orthologs in the other two lineages as a proxy for the ancestral state.

We first looked for evidence that orthologs of X–Y gene pairs from therian mammals are enriched for expression in testis. We calculated testis specificity in chicken using a panel of 15 adult male tissues (McCarthy et al. 2019) and in five-pacer viper using a panel of four adult male tissues (Yin et al. 2016). In each case, after controlling for the effects of broad expression, expression in testis had no significant effect on the odds of ancestral X genes retaining a surviving Y homolog (Fig. 4).

Likewise we searched for evidence that orthologs of Z–W gene pairs from birds were ancestrally enriched for expression in ovary. We calculated ovary specificity in human using a panel of 39 adult female tissues (GTEx Consortium 2017) and in eastern garter snake using a panel of eight adult female tissues (Perry et al. 2018). We did not detect a significant effect of ovary-specific expression in either lineage on the survival of W-linked genes in birds (Fig. 4).

Lastly, using the human (GTEx Consortium 2017) and chicken (McCarthy et al. 2019) ovary specificities, we found that ovary-specific expression in either lineage had no effect on the survival of W-linked genes in snakes (Fig. 4). We conclude that ancestral bias for expression in sex-specific tissues played no significant role in survival of X–Y and Z–W pairs.

### Survivors are not significantly enriched for genes with conserved sex bias in mammals

We also asked whether survivors were enriched for genes that show a conserved sex bias in mammals (Naqvi et al. 2019). Because surviving X–Y gene pairs preserve ancestral gene expression levels in both sexes, the X-homologs of X–Y gene pairs are not subject to X-inactivation, and are expressed more highly from two copies in females than they are from a single copy in males. As a result, the X-homologs of X–Y gene pairs were called as having a conserved female bias in this dataset; therefore we excluded therian sex-linked genes from this analysis. Since both birds and snakes have female-specific W chromosomes, we looked for an enrichment of genes with a conserved female bias in one or more tissues in the mammalian autosomal orthologs of surviving Z–W gene pairs, relative to other ancestral genes (Table 8). We did not observe an enrichment for conserved female-biased genes: while 54 of 275 survivors were female-biased, so were 298 of 1658 non-survivors (Table 8).

**Table 8.** Survival of conserved female-biased genes in mammals.

|  | Conserved Female Biased | Other Genes |
| --- | --- | --- |
| Z-W Pairs | 54 | 221 |
| Other Ancestral Z | 298 | 1360 |

### Dosage sensitivity and expression breadth made independent contributions to survival

We sought to explore the interactions between dosage sensitivity, breadth of expression, and strength of purifying selection in the survival of genes on sex-specific chromosomes in our combined data set. Therefore, we set out to quantify the survival of each ancestral gene. Within a species, the sex-specific homologs of ancestral gene pairs were exposed to the forces of genetic decay for different periods of time, depending on the age of their evolutionary strata. We looked for evidence of evolutionary strata in caenophidian snakes (Supplemental Figs. S1-S3, and S7, Supplemental Tables S2 and S8, Methods) and tabulated published information about evolutionary strata in birds and mammals (Supplemental Fig. S1, Supplemental Tables S9 and S10). We calculated a gene-wise ‘survival fraction’ to control for the different numbers of sampled species and evolutionary strata of different ages when comparing across X–Y and Z–W systems. For each gene within a lineage, we calculated its total branch length across the tree of species, then divided this by the maximum branch length possible for a gene of its evolutionary stratum (Supplemental Tables S8-S10, Methods). These values range from 0 (lost in all lineages) to 1 (survival in every possible lineage) (Supplemental Table S5).

We used principal components analysis to explore the influence of factors related to dosage sensitivity, breadth of expression, and strength of purifying selection on the survival of ancestral gene pairs across all three lineages (Fig. 5). The first principal component separated genes according to their survival fraction, allowing us to visualize the contribution of each factor to survival. Consistent with the hypothesis that intense purifying selection maintains surviving gene pairs, the dN/dS ratios in mammals, birds, and reptiles were most closely aligned with the survival fraction (although they point in the opposite direction, as survivors have a reduced dN/dS ratio). Human haploinsufficiency probabilities are closely aligned with survival; these probabilities reflect the output of a model trained on known human haplosufficient genes that takes multiple factors as input, including embryonic expression levels and strength of purifying selection (Huang et al. 2010). Measures of dosage sensitivity and breadth of expression make distinct and approximately orthogonal contributions to the survival of genes on sex-specific chromosomes. This suggests that some genes may survive primarily due to dosage sensitivity, while others may be preserved mostly because they are required in many tissues and cell types.

**Figure 5.**
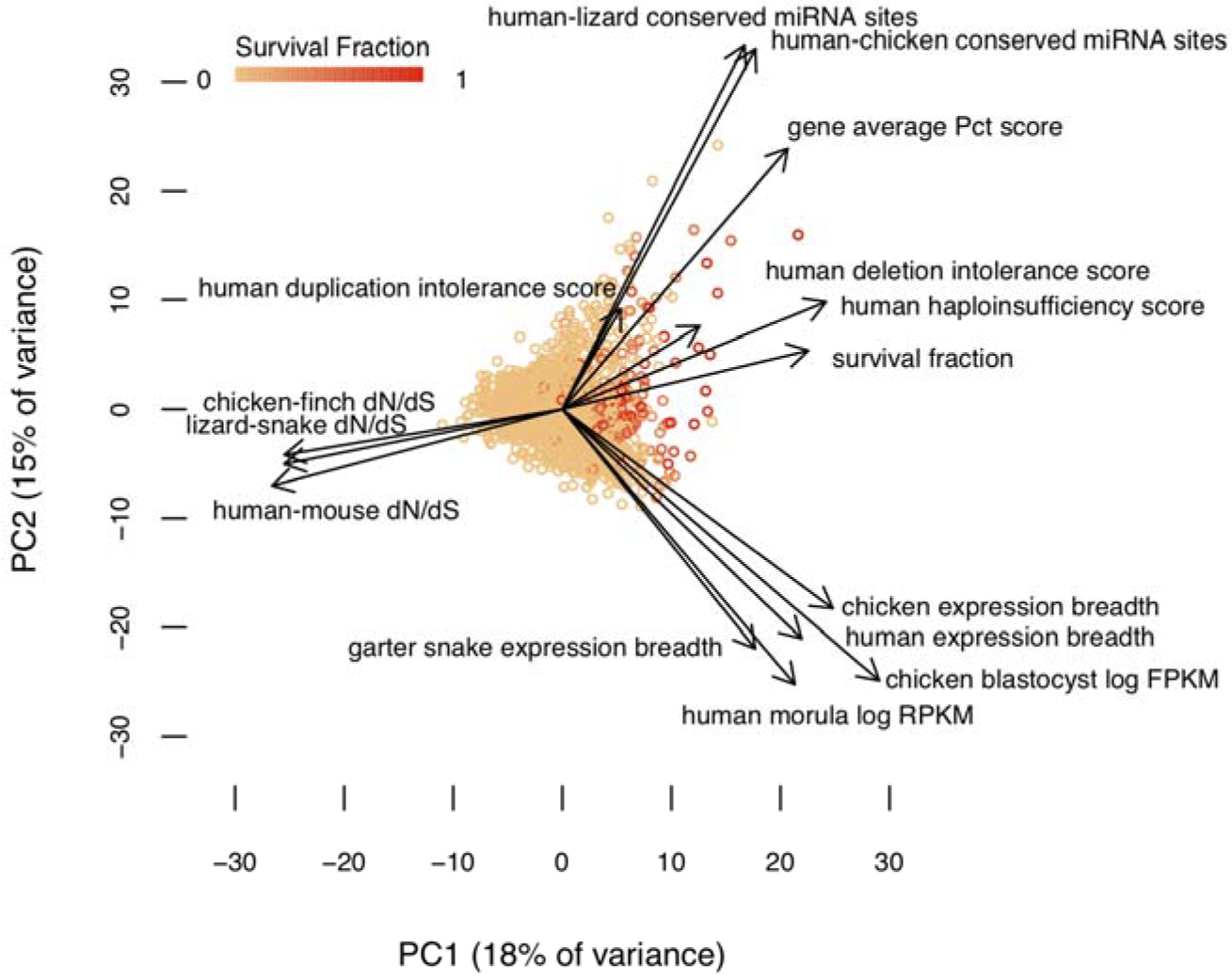
Dosage sensitivity and broad expression make independent contributions to survival. A statistical summary of survival factors from 2573 genes based on principal component axis one (PC1) and axis two (PC2). Points represent individual genes, colored by their survival fraction from orange (no survival) to red (survival in all possible lineages) (See Methods and Supplemental Table S5). Arrows show the contribution of each factor to the variation in survival of ancestral genes on sex-specific chromosomes. Dosage sensitivity and breadth of expression make roughly orthogonal contributions, while strength of purifying selection is closely aligned with survival.

## Discussion

We were unable to detect a strong influence of sex and reproduction related functions in the survival of ancestral genes across sex-specific chromosomes in amniotes. If genes in these functional categories are not highly dosage sensitive, broadly expressed, and subject to intense purifying selection, then this may hinder their survival on sex-specific chromosomes. Genes involved in sex determination and gonadogenesis may be poorly represented among survivors because they are less dosage sensitive. In vertebrates, development of the bipotential gonads is canalized down two pathways to generate either ovaries or testes; the regulatory cascades that govern this decision are a series of bistable switches, robust to perturbation (Munger and Capel 2012). As a consequence, the genes and networks regulating gonadal sex determination may be less affected by subtle changes in gene dose than those of developmental regulators that produce a graded response with changes in gene dose, like morphogens. Likewise, genes with expression limited to sex-specific tissues and genes with sex-biased expression are both under reduced selective constraint, which could reduce their chances of resisting genetic decay. In contrast, genes expressed across many genes and tissues are subject to complex selection pressures that result in more intense purifying selection (Duret and Mouchiroud 2000) and greater conservation of patterns of gene expression (Yang et al. 2005). Sex-biased genes are more narrowly expressed than genes with no bias in *Drosophila* (Meisel 2011), birds (Mank et al. 2008) and mammals (Mank et al. 2008; Meisel 2011; Naqvi et al. 2019), and this narrow expression is accompanied by reduced selective constraint (Meisel 2011; Naqvi et al. 2019). Taken together, this suggests that the selective pressures that enriched surviving Z–W and X–Y pairs for broadly expressed, dosage sensitive regulators under strong purifying selection may present obstacles to the survival of sex and reproduction related genes.

Selection for sex and reproduction related functions may be more influential at specific stages of sex chromosome evolution, making them more difficult to detect in our analyses of amniote sex chromosomes. The sex chromosomes of caenophidian snakes, birds, and therian mammals are all evolutionarily old and highly differentiated. As a result, our analyses will not capture the influence of factors related to sex and reproduction on gene survival during the early stages of sex chromosome evolution. In *Drosophila miranda*, surviving ancestral genes on Y chromosomes are enriched for genes with ancestral male-specific expression, but this effect is strongest among the 5% of ancestral genes (109 / 2250 ancestral genes) that have been retained on the Y chromosome, but lost from the X chromosome (Kaiser and Bachtrog 2014). There is no parallel to this high rate of gene loss on amniote Z and X chromosomes, so these observations may reflect selective pressures unique to the *Drosophila* lineage, rather than a common process in the early stages of sex chromosome evolution.

Selection for sex and reproduction related functions could also lead to changes in the function or expression patterns of ancestral genes later in sex chromosome evolution. We previously reported that ancestral Y-linked genes that were amplified into multi-copy gene families in at least one species survived through more of therian Y chromosome evolution than those that remained single copy, and we noted that these multi-copy families were expressed exclusively or predominantly in the testis (Bellott et al. 2014). While the evolution of testis-specific expression likely preceded gene amplification on the Y chromosome, most of these genes were ancestrally broadly expressed. Because our analyses here focus on the ancestral expression patterns of survivors and non-survivors, they cannot capture changes in expression pattern that occur during the evolution of sex-specific chromosomes, and as a result they may underestimate the influence of selection for sex and reproduction related functions. However, specialization for reproductive function does not inevitably occur on sex-specific chromosomes, even for multi-copy gene families. All of the genes on the female-specific W chromosome of chicken, including the multi-copy *HINTW*, retain their ancestral broad expression (Bellott et al. 2017). Studies of additional avian and caenophidian W chromosomes will reveal how frequently ancestral genes evolve expression restricted to female reproductive tissues.

Like their counterparts on mammalian Y chromosomes (Bellott et al. 2014) and avian W chromosomes (Bellott et al. 2017), the surviving ancestral genes on caenophidian W chromosomes are enriched for broadly expressed, dosage sensitive regulators under strong purifying selection. Our findings across these three highly differentiated sets of amniote sex chromosomes have parallels in the non-random survival of ancestral genes on the evolutionarily young Y chromosomes of *Drosophila miranda* (about 1.5 MY) (Kaiser et al. 2011) and threespine stickleback (less than 16 MY)(White et al. 2015). Surviving gene pairs on the Neo-X and Neo-Y chromosomes of *Drosophila miranda* are expressed at higher levels and across more tissues than genes lost to decay (Kaiser et al. 2011). An abundance of protein-protein interactions is correlated with dosage sensitivity (Papp et al. 2003; Huang et al. 2010); in both threespine stickleback and *Drosophila miranda*, surviving X–Y gene pairs were enriched for genes encoding proteins with many partners in protein-protein interaction networks (White et al. 2015; Kaiser et al. 2011). The repeated finding that sex-specific chromosomes preferentially retain broadly expressed, dosage-sensitive genes, regardless of whether the chromosomes are female-specific or male-specific, or in the early or late stages of differentiation, demonstrates that purifying selection is effective at preserving the correct dosage of critical ancestral genes in the absence of crossing over. We propose that the survival of ancestral genes on sex-specific W and Y chromosomes is driven by pressure to maintain the viability of the heterogametic sex.

One corollary of this proposal is that there will be severe consequences for monosomy of sex chromosomes that carry an abundance of dosage sensitive Z–W or X–Y gene pairs. We previously proposed that the surviving X–Y gene pairs in humans and Z–W gene pairs in chicken were essential for the viability of heterogametic embryos (Bellott et al. 2014, 2017). In humans, 99% of human 45,X conceptuses are inviable, while the remainder are often mosaic for all or part of a second sex chromosome (Hook and Warburton 1983; Hassold et al. 1988; Cockwell et al. 1991). Similarly, sex chromosome monosomy has been observed in chicken embryos at the blastocyst stage (Fechheimer 1981), but these embryos do not survive beyond five days of incubation (Fechheimer 1981; Bloom 1972). Despite nearly six decades of cytogenetic observations of the caenophidian sex chromosomes (Kobel 1962; Becak et al. 1962), including an unusual 3A:ZZW intersex twin-spotted ratsnake (*Elaphe bimaculata*)(Rovatsos et al. 2018), we are unaware of any reports of sex chromosome monosomy in a caenophidian snake. Given the number of dosage sensitive genes among the surviving ancestral genes on caenophidian W chromosomes, as well as their ancestrally broad expression, extending to early embryogenesis, we predict that sex chromosome monosomy is also embryonic lethal in caenophidian snakes.

Dosage compensation on Z and X chromosomes also functions to maintain the viability of the heterogametic sex and is so deeply entwined with the survival of ancestral genes on W and Y chromosomes that they cannot be studied independently. Surviving Z–W and X–Y gene pairs maintain expression in the heterogametic sex at levels similar to those of pseudoautosomal or autosomal orthologs for the same sex in close outgroups, but not necessarily equal expression between the homogametic and heterogametic sex. Depending on its intrinsic sensitivity to changes in expression level, each gene may or may not evolve dosage compensation to restore this ancestral expression level after the decay of a sex-specific W or Y-linked homolog (Naqvi et al. 2018). In genes that are more sensitive to underexpression than overexpression, autosomal paralogs derived from ancestral sex-linked genes can rescue loss of sex-specific homologs (Hughes et al. 2015). To fully account for these factors, a comprehensive characterization of the nature and extent of dosage compensation in caenophidian snakes will require the development of comparable gene expression data from a female caenophidian snake and henophidian or lizard outgroup species, paired with highly contiguous, reference-grade sequences of both Z and W sex chromosomes as well as the autosomes.

Our analyses revealed that dosage sensitivity and breadth of expression contribute independently to survival of genes on sex-specific chromosomes. This suggests that genes expressed across a broad array of tissues face complex selective pressures that make an important contribution to survival, distinct from increasing sensitivity to changes in gene dose. One possible explanation is that pleiotropic effects may impose constraints on the evolution of dosage compensation for broadly expressed genes. Models of the evolution of sex-biased gene expression predict that genes preferentially expressed in the heterogametic sex are less likely to evolve on the Z and X chromosomes than on autosomes when they are subject to pleiotropic constraints (Connallon and Clark 2010). Ancestral genes expressed across a broad array of tissues are therefore less likely to evolve increased expression from their Z or X homologs than those expressed more narrowly. As a result, mutations that decrease expression from the sex-specific homolog will more frequently arise opposite a Z-linked or X-linked homolog that cannot compensate for their reduced dosage, increasing the stringency of selection against their loss. Under this model, even with equivalent levels of dosage-sensitivity, more broadly-expressed genes should prove more resistant to genetic decay than those that are more narrowly expressed.

Summed across all three lineages, nearly an eighth of the ancestral amniote genome was subjected to 2.5 billion years of sex chromosome evolution, winnowing 2573 ancestral genes to just 322 survivors on sex-specific chromosomes. These survivors are enriched for global regulators of gene activity, with broad expression across tissues and developmental time. Homologs of survivors play key roles in human biology. Previously we reported that human orthologs of survivors in birds are enriched for genes implicated in human disorders attributed to haploinsufficiency (Bellott et al. 2017), and that several X-linked intellectual disability syndromes have been mapped to the X-linked homologs of mammalian X-Y pairs (Bellott et al. 2014). Here we extend those findings to show that the human orthologs of survivors regulate a broad range of processes in early embryonic development across multiple organ systems. Because the human homologs of survivors are depleted for variation in copy number and coding sequence, we anticipate that even subtle modulation of the transcription, splicing, translation, and stability of survivors will have disproportionately large effects on development and disease.

## Methods

### Reconstructing ancestral sex chromosomes

We downloaded peptide sequences and lists of human, chicken, and lizard orthologs from Ensembl (release 94)(Zerbino et al. 2018). To identify snake orthologs, we aligned these peptides to the reference genome assemblies of prairie rattlesnake (GCA_003400415.2)(Schield et al. 2019), mainland tiger snake (GCA_900518725.1), five-pacer viper (Deinagkistrodon_acutus.fa.gz)(Yin et al. 2016), and *Boa constrictor* (snake_6C_scaffolds.fa.gz)(Bradnam et al. 2013), using blat (Kent 2002) in tblatx mode (-t=dnax –q=prot), and retained only the best hit for each peptide sequence.

We seeded our search for the ancestral set of sex-linked genes with 106 genes previously mapped to the Z or W chromosomes of caenophidian snakes (Matsubara et al. 2012, 2016; Vicoso et al. 2013a; Rovatsos et al. 2015; Yin et al. 2016; Perry et al. 2018) and the orthologous python Y chromosome (Gamble et al. 2017) (Supplemental Table S1). Most of these genes have orthologs on human chromosomes 3, 7, 10, 17, and 19; chicken chromosomes 2 and 27; and lizard chromosome 6.

We identified genes as sex-linked in the ancestor of caenophidian snakes by looking for orthologs of human, chicken, and lizard genes that were syntenic with published sex-linked genes in one or more of the three caenophidian snake genome assemblies, starting with prairie rattlesnake (the most contiguous). We used orthologous scaffolds in the *Boa constrictor* assembly as an outgroup to resolve whether lineage-specific gains and losses occurred before or after the divergence of henophidian and caenophidian snakes.

We found that 99 of 106 published snake sex-linked genes mapped to the prairie rattlesnake Z chromosome assembly (CM012323.1). Of the seven remaining genes, five (*ARF1, KLF6, MRPL3, MYO1D*, and *ZBTB47*) mapped to autosomes in prairie rattlesnake, but were found on scaffolds syntenic with other Z-linked genes in tiger snake and five-pacer viper, and on autosomes orthologous to the Z chromosome in outgroup species. The other two (*PINX1, TMEM35B*) are not syntenic with caenophidian Z-linked genes or their orthologs in any species, and probably represent genes acquired by the python Y chromosome (Supplemental Table S2).

We identified 1652 genes as ancestral to the caenophidian sex chromosomes, including 1500 genes from the prairie rattlesnake Z chromosome assembly, and 152 others based on synteny in other species (Supplemental Table S2). This set of ancestral genes includes 80 published W-linked genes from 13 caenophidian snake species (Matsubara et al. 2006, 2016; Vicoso et al. 2013a; Yin et al. 2016; Perry et al. 2018)(Supplemental Table S3). We also identified another 41 genes among our ancestral set that were previously identified as candidate W-linked genes in pygmy rattlesnake and mountain garter snake (Vicoso et al. 2013a), but were rejected because their lizard orthologs did not map to lizard chromosome 6 (Supplemental Table S4).

Our ancestral set of genes contains 348 members of 8 multi-copy gene families, including 18 of 41 candidate W-linked genes (Supplemental Table S2). Several of these gene families, particularly the olfactory receptor, vomeronasal receptor, and immunoglobulin heavy chain variable region families, form megabase-sized arrays that likely vary in copy-number between species. We excluded genes from 8 multi-copy families and restricted our analyses to single-copy genes where we could confidently assign orthologs across species, yielding a set of 1304 ancestral genes (Supplemental Table S2).

### Reconstructing Evolutionary Strata

#### Identifying homologs of ancestral genes on the caenophidian sex chromosomes for phylogenetic analysis

We seeded our search for homologs of ancestral genes with coding sequences of the longest isoforms of human, chicken, and lizard orthologs from Ensembl. To expand our resolution in the caenophidian lineage, we used exonerate (version 2.2.0)(Slater and Birney 2005) to predict coding sequences from genomic reference sequences in *Boa constrictor (*Bradnam et al. 2013), Burmese python (*Python molurus bivittatus*) (GCF_000186305.1)(Castoe et al. 2013), speckled rattlesnake (*Crotalus mitchellii*) (GCA_000737285.1)(Gilbert et al. 2014), prairie rattlesnake (Schield et al. 2019), timber rattlesnake (*Crotalus horridus*) (GCA_001625485.1)(Sanders, et al. Unpublished), pygmy rattlesnake (Vicoso et al. 2013a), Taiwan habu (*Protobothrops mucrosquamatus*) (GCF_001527695.2)(Aird et al. 2017), Okinawa habu (*Protobothrops flavoviridis*) (GCA_003402635.1)(Shibata et al. 2018), five-pacer viper, adder (*Vipera berus berus*) (GCA_000800605.1)(Liu, et al. Unpublished), mountain garter snake (Vicoso et al. 2013a), eastern garter snake (*Thamnophis sirtalis*) (GCF_001077635.1)(Perry et al. 2018), corn snake (*Pantherophis guttatus*) (GCA_001185365.1)(Ullate-Agote et al. 2014), mainland tiger snake, and king cobra (*Ophiophagus hannah*) (GCA_000516915.1)(Vonk et al. 2013) (Supplemental Data S1).

#### Phylogenetic analyses

For cross-species phylogenetic analyses, we generated multiple alignments using Clustal Omega (version 1.2.4)(Sievers et al. 2011) and PAL2NAL (version 14)(Suyama et al. 2006) (Supplemental Data S2). For each alignment, we generated a set of phylogenetic trees based on the species trees from TimeTree (Kumar et al. 2017) (Supplemental Fig. S1), allowing for stratum formation events in the common ancestor, or in later lineages. We used TreePuzzle (version 5.2)(Schmidt et al. 2002) to estimate the gamma distribution parameter and transition-transversion ratio. We used TreePuzzle together with DNAML and DNAPARS in PHYLIP (version version 3.696)(Felsenstein 1989) to evaluate which trees were most consistent with the multiple alignment, using the Shimodaira-Hasegawa test (Shimodaira and Hasegawa 1999).

#### Pseudoautosomal Boundaries

To locate the pseudoautosomal boundaries in snakes, we mapped male and female reads of pygmy rattlesnake (PRJNA210004)(Vicoso et al. 2013a) to the prairie rattlesnake genome, and male and female reads from mountain garter snake (PRJNA210004)(Vicoso et al. 2013a) to the eastern garter snake genome (*Thamnophis sirtalis*)(GCF_001077635.1)(Perry et al. 2018) using bowtie2 (version 2.3.4.1)(Langmead et al. 2009). We aligned the eastern garter snake genome with the prairie rattlesnake genome using LAST (version 992)(Kielbasa et al. 2011) to generate a chain file, which we filtered with chainSort, chainNet, and netChainSubset from the UCSC genome browser tools (version 0.0.3-1)(Haeussler et al. 2019). We converted read alignments into a coverage bedGraph using bedtools (version 2.26.0)(Quinlan and Hall 2010), and then used liftOver from the UCSC genome browser tools to map coordinates from eastern garter snake to the more contiguous prairie rattlesnake assembly. We plotted the base 2 logarithm of the ratio of normalized female to normalized male coverage across the Z chromosome in 100 kilobase windows, to visualize the transition from equal read depth in both sexes in the pseudoautosomal region to lower coverage in females in the younger evolutionary strata on the Z chromosome. We note that read coverage in ancestral arrays of multicopy gene families was greater in mountain garter snake females than in males, but we cannot distinguish between extreme polymorphism in array size on the Z chromosome between individuals, or expansion of these arrays on the W chromosome.

#### Caenophidian Evolutionary Strata

Phylogenetic analyses of ZW gene pairs reveal that mountain garter snake, five-pacer viper and pygmy rattlesnake share a common evolutionary stratum, containing 981 ancestral single-copy genes. This stratum formed after these three species diverged from the Arafura file snake, but before they diverged from each other, between 76 and 61.7 MYA (Fig. 1A, Supplemental Figs. S1 and S7, Supplemental Tables S2 and S8). We infer that one or more parallel strata including *WAC* and *CTNNB1* formed on the lineage leading to arafura file snake (Supplemental Figs. S1 and S7).

The most distal Z homologs of W-linked genes in this old stratum lie near an array of olfactory receptor genes (Supplemental Table S2). Similar arrays exist on the orthologous autosomes in chicken and human, indicating that this array is an ancestral feature of the amniote genome. This array is not fully resolved in the green anole lizard genome, and the lizard orthologs of more distal Z-linked genes are in unassigned scaffolds. Because W-linked genes were only predicted for orthologs of lizard chromosome 6 (Yin et al 2016), *SNRNP70* from this stratum is the most distal Z–W pair predicted for five-pacer viper. However, the pseudoautosomal boundary of five-pacer viper maps to a much more distal location in prairie rattlesnake (Schield et al. 2019), so we infer that one or more additional strata must have formed in the lineage leading to five-pacer viper (Supplemental Fig. S1).

In mountain garter snake, 64 ancestral single-copy genes including *SRRM2* and *PPP6R1* belong to a stratum that formed in the common ancestor of corn snake and mainland tiger snake (Supplemental Figs. S1 and S2, Supplemental Tables S2 and S8), around 51 MYA. Subsequently, around 48 MYA, a stratum containing 197 ancestral single-copy genes, including *PPP6R1* and *PRF1*, formed in the ancestor of mountain garter snake and corn snake (Supplemental Figs. S1 and S2, Supplemental Table S2). The remaining 62 single-copy genes map to the opposite side of an array of vomeronasal receptor genes and appear to be pseudoautosomal (Supplemental Table S2, Supplemental Figs. S2 and S8). A parallel stratum, including *PPP6R1*, formed independently in the lineage leading to mainland tiger snake, but we could not resolve whether this predates the divergence with king cobra (Supplemental Figs. S1 and S2).

A stratum containing 216 genes, including *RASIP1* and *RPL18*, formed in the common ancestor of the pygmy rattlesnake and snakes from the *Crotalus* and *Protobothrops* genera (Supplemental Figs. S1 and S3, Supplemental Tables S2 and S8), around 29.5 MYA. This was followed by a stratum containing 45 genes, including *FLT3LG*, in the common ancestor of pygmy rattlesnake and timber rattlesnake (Supplemental Fig. S3, Supplemental Table S2), about 12.5 MYA. We infer that a parallel stratum containing *FLT3LG* formed in the common ancestor of Taiwan habu and Okinawa habu (Supplemental Figs. S1 and S3). Pygmy rattlesnake shares a common pseudoautosomal region boundary with the five-pacer viper and the prairie rattlesnake (Schield et al. 2019), near the same array of vomeronasal receptor genes as in garter snake. This suggests that this array of genes may participate in recurrent structural rearrangements that initiate stratum formation (Supplemental Fig. S3, Supplemental Tables S2 and S8).

#### Avian Evolutionary Strata

We reconciled published stratum assignments in birds (Handley et al. 2004; Vicoso et al. 2013b; Wright et al. 2014; Zhou et al. 2014; Xu et al. 2019a; Wang et al. 2019) with our new reconstruction of the ancestral gene order on the avian Z chromosome (Supplemental Table S9). We used ANGES (version 1.01) (Jones et al. 2012) to reconstruct the order of chicken orthologs on the ancestral autosome that evolved into the avian Z and W sex chromosomes. Briefly, we aligned the peptide sequences of chicken Z-linked genes from Ensembl (release 94)(Zerbino et al. 2018) to the reference genome assemblies of chicken (GCF_000002315.6)(Hillier et al. 2004; Bellott et al. 2010), turkey (*Meleagris gallopavo*) (GCF_000146605.3)(Dalloul et al. 2010), duck (*Anas platyrhynchos*) (GCF_003850225.1)(Zhang et al. 2014), Anna’s hummingbird (*Calypte anna*) (GCF_003957555.1)(Zhang et al. 2014), great tit (*Parus major*) (GCF_001522545.3)(Laine et al. 2016), common canary (*Serinus canaria*) (GCF_007115625.1)(Frankl-Vilches et al. 2015), dark-eyed junco (*Junco hyemalis*) (GCA_003829775.1)(Friis et al. 2018), ostrich (*Struthio camelus australis*) (GCF_000698965.1)(Zhang et al. 2014), cassowary (*Casuarius casuarius*) (GCA_003342895.1)(Sackton et al. 2019), emu (*Dromaius novaehollandiae*) (GCF_003342905.1)(Sackton et al. 2019), white-throated tinamou (*Tinamus guttatus*) (GCF_000705375.1)(Zhang et al. 2014), and Chilean tinamou (*Nothoprocta perdicaria*) (GCF_003342845.1)(Sackton et al. 2019)], as well as the assemblies of American alligator (*Alligator mississippiensis*) (GCF_000281125.3)(St John et al. 2012), green anole lizard (GCF_000090745.1)(Alföldi et al. 2011), and human (GCF_000001405.26)(International Human Genome Sequencing Consortium 2004) genomes as outgroups. We recovered seven contiguous ancestral regions in the ancestor of birds. The largest contiguous ancestral region contains 472 genes and stretches from the boundary between stratum 0 and stratum 1 to the pseudoautosomal region, and provides sufficient information to reconstruct all lineage-specific strata in birds (Supplemental Table S5). All other contiguous ancestral regions reside within stratum 0, which formed in the common ancestor of all birds and has been subject to extensive lineage-specific rearrangments on the Z chromosome (Zhou et al. 2014). We made our best effort to reconstruct the ancestral gene order based on outgroup species within stratum 0, but our downstream analyses do not depend on the exact order of genes within this stratum.

#### Mammalian Evolutionary Strata

In mammals, we relied on published Y chromosome sequence assemblies and reconstructions of evolutionary strata (Supplemental Table S10) (Lahn and Page 1999; Skaletsky et al. 2003; Ferrante et al. 2003; Murphy et al. 2007; Van Laere et al. 2008; Chang et al. 2011; Li et al. 2013; Bellott et al. 2014; Skinner et al. 2016; Janečka et al. 2018).

### Functional annotation

We mapped published functional annotations onto our set of ancestral genes and their human orthologs. For expression breadth, we normalized the expression of each gene to the highest expression in any tissue, and took the average expression across all tissues. We used UniProt annotations to identify caenophidian Z–W pair genes involved in regulatory processes.

### OMIM

We downloaded the full text of OMIM (McKusick-Nathans Institute of Genetic Medicine 2019) and searched entries of the human orthologs of ancestral caenophidian Z-linked genes for “haploinsufficient” or “haploinsufficiency”, limiting our search to phenotypes with a known molecular basis. We examined each of the resulting entries to verify that there was evidence that the phenotype was caused by haploinsufficiency.

### Gene Ontology

We used the PANTHER statistical overrepresentation test (Mi et al. 2019a; Mi et al. 2019b) using Fisher’s exact test with false discovery rate correction to identify functional annotations that were enriched among the human orthologs of surviving Z–W and X–Y pair genes, relative to the reference list of orthologs of other ancestral Z and X genes. We selected the orthologs of ancestral Z and X genes as a reference list, instead of all human genes, to control for any functional coherence among ancestral genes that pre-dated the start of genetic decay on W or Y chromosomes.

### Logistic regression

Logistic regression models of survival were constructed using the function “glm” in the `stats` package of R 3.6.1, with the option `family = binomial`. For each species, we used kallisto and sleuth to estimate TPM values across male or female tissues. We calculated the tissue specificity index as the fraction of TPMs originating from testis or ovary across a panel of male or female tissues, respectively. We calculated expression breadth as previously described (Bellott et al. 2014).

### Survival Fraction

To calculate longevity, we summed all branch lengths in the most parsimonious tree from each of the species where a gene is present to the last common ancestor before stratum formation. We divided the longevity for each gene by the maximum possible longevity for each stratum to generate the survival fraction. Survival fractions range from 0 (lost in all lineages) to 1 (retained in each lineage). In cases where a gene was transposed from the X chromosome to the Y chromosome after the ancestral Y gene was lost, we treated the transposition as an additional stratum.

### Principal components analysis

We modeled the survival fraction as a linear combination of the functional annotations for each gene using the `prcomp` function in the `stats` package of R 3.6.1, with options `scale = TRUE, center = TRUE, na.action=na.aggregate` to normalize the annotation data and replace missing values with the mean value for each annotation. We used the fourteen quantitative annotation categories we identified as enriched among survivors on sex-specific chromosomes across amniotes: human haploinsufficiency score, human deletion intolerance score, human duplication intolerance score, gene average Pct score, the number of human-chicken conserved miRNA sites, the number of human-lizard conserved miRNA sites, human expression breadth, chicken expression breadth, garter snake expression breadth, human morula mRNA expression levels, chicken blastocyst mRNA expression levels, human-mouse dN/dS, chicken-finch dN/dS, and lizard-snake dN/dS.

### Statistics

Details of all statistical tests (type of test, test statistic, and P-value) used in this article are provided in Supplemental Table S11.

## Data Access

Data supporting the findings of this study are available within the paper and its Supplemental information files.

## Acknowledgements

We thank R. Edwards, T. Amos, J. Tang, B. Cawood, S. Rispin, D. E. Tuipulotu, and P. Waters for permission to analyze data from their unpublished assembly of the mainland tiger snake genome; W. S. Sanders, M. A. Arick Jr. A. Thrash, D. Rhoads, J. Pummill, S. Beaupre, B. Nanduri, A. Perkins, D. G. Peterson, and R. D. Steele for permission to analyze data from their unpublished assembly of the timber rattlesnake genome; Y. Liu, D. Hughes, H. Dinh, S. Dugan, S. Jhangiani, S. Lee, G. Okwuonu, J. Santibanez, D. Bandaranaike, H. Chao, M. Dahdouli, R. Gnanaolivu, S. Gross, W. Hale, M. Javaid, J. Jones, S. Matakis, M. Munidasa, A. Narasimhan, D. Ngo, L. Nguyen, E. Skinner, E. Ongeri, N. Osuji, R. Puazo, S. Qi, C. Qu, J. Quiroz, R. Raj, K. Rajbhandari, X. Yun, Y. Han, C. Kovar, J. Reid, H. Doddapaneni, T. Madsen, B. Ujvari, D. Muzny, K. Worley, and R. Gibbs for permission to analyze data from their unpublished assembly of the European adder genome; A. Godfrey and S. Naqvi for assistance with the analysis of GTEx data; and helpful discussions and A. Godfrey, E. Jackson, J. Hughes and H. Skaletsky for critical reading of the manuscript. This work was supported by the Whitehead Institute, the Howard Hughes Medical Institute, and generous gifts from Brit and Alexander d’Arbeloff and Arthur W. and Carol Tobin Brill.

## Author Contributions

D.W.B. and D.C.P. designed the study and wrote the paper. D.W.B. performed the analyses.

## Disclosure Declaration

The authors declare no competing financial interests.

